# Cell type and subcellular compartment specific APEX2 proximity labeling proteomics in the mouse brain

**DOI:** 10.1101/2021.04.08.439091

**Authors:** V. Dumrongprechachan, G. Soto, M.L. MacDonald, Y. Kozorovitskiy

**Affiliations:** Department of Neurobiology, Northwestern University, Evanston, IL 60208; The Chemistry of Life Processes Institute, Northwestern University, Evanston, IL 60208; Department of Psychiatry, University of Pittsburgh, Pittsburgh, PA 15213; Biomedical Mass Spectrometry Center, University of Pittsburgh, Pittsburgh, PA 15213

## Abstract

The vertebrate brain consists of diverse neuronal types, classified by distinct anatomy and function, along with divergent transcriptomes and proteomes. Defining the cell type-specific neuroproteome is important for understanding the development and functional organization of neural circuits. This task remains challenging in complex tissue, due to suboptimal protein isolation techniques that often result in loss of cell-type specific information and incomplete capture of subcellular compartments. Here, we develop a genetically targeted proximity labeling approach to identify cell-type specific subcellular proteome in the mouse brain. Using adeno-associated viral transduction, we express subcellular-localized APEX2 to map the proteome of the nucleus, cytosol, and cell membrane of Drd1 receptor-positive striatal neurons. We show that each APEX2 construct can differentially and rapidly biotinylate proteins *in situ* across various subcellular compartments, confirmed by imaging, electron microscopy, and mass spectrometry. This method enables flexible, cell-type specific quantitative profiling of subcellular proteome in the mouse brain.

## Main

The central nervous system (CNS) is made up of functionally and morphologically diverse types of neurons defined by their proteome and transcriptome^1^. Different classes of neurons throughout the brain form millions of intertwined short and long-range synaptic connections to mediate and regulate neurotransmission, controlling sensory processing and behavior. Recent single-cell RNA sequencing (scRNA-seq) of the rodent brain revealed a vast array of CNS-region specific neuronal types and subtypes, enabling an unbiased mapping of molecular identity and functional anatomy^2^. Although scRNA-seq provides cell-type specific information about transcription, scRNA-seq data lack information about protein subcellular localization, which influences protein function. Because proteins are the ultimate product of gene expression, the effectors that maintain and regulate cellular physiology, proteomic analyses yield a direct readout of cellular identities and states. Therefore, mapping the proteome with both cell-type and compartment specificity is crucial for understanding the coordinated functions of neural circuits in the vertebrate brain.

In many model systems, identification of distinct cell classes in the brain is achieved by conditional expression of site-specific recombinases and fluorescent reporters, or other effectors, under the control of a gene-specific promoter^3^. Cell isolation approaches such as tissue dissociation, followed by fluorescent cell sorting (FACS)^4^ or laser capture microdissection (LCM)^5^, are usually employed to isolate reporter-positive somata for proteomics analyses. Whole-cell patch clamp can also be used to collect cytoplasmic content for single-cell assays^6^. In addition, biochemical fractionation methods are often used to access microdomain-specific proteomes such as nuclear, membrane, and synaptosomal fractions^7^. Such preparations typically do not yield cell-type specific data. Despite improvements in protein isolation and biochemical fractionation techniques, these methods are not ideal for mapping cell-type specific neuronal proteomes due to a substantial loss of peripheral cellular structures such as dendrites and axons, containing physiologically important information.

Recent advances in bio-orthogonal strategies for neuroproteomics facilitated the capture of proteins in a cell type-specific manner. Genetically encoded engineered tRNA synthetase and/or tRNA pair were expressed to metabolically incorporate non-canonical amino acids (NCAAs) into the proteome of a specific cell type (*e*.*g*., MetRS for methionine^8^, SORT_CAU_ for tyrosine^9^). Subsequently, cell-wide labeled proteins were biotinylated via click chemistry and enriched for identification and quantification^8,9^. Because incorporation of NCAAs depends on protein translation, this approach is not suitable for answering questions about many rapid biological processes. The experimental time window of NCAA incorporation can take up to several days^8,9^.

Proximity labeling (PL) is an alternative approach to bio-orthogonal labeling. Genetically encoded labeling enzymes such as BioID^10^, TurboID^11^, and APEX2^12^ can be expressed and localized to a specific subcellular compartment. In the presence of exogenous biotin substrates, *in situ* biotinylation occurs rapidly, from minutes to hours for TurboID, and within seconds for APEX2. This technique enables taking the snapshots of the proteome with restricted spatial labeling and on short temporal timescales. Therefore, PL methods offer both cell-type and subcellular compartment specific information about the identified proteins. Prior studies demonstrated the effectiveness of PL approaches for proteomic mapping in many model organisms such as *D. melanogaster*^13^. Transgenic mouse model is a prevalent model in neuroscience; yet, there were only a few studies using BioID directly in the mouse brain tissues^14,15^, while HRP/APEX2-based proteomics, which offers superior labeling speed, were previously used in primary neuronal cultures alone^16,17^.

Here, we described a genetically targeted APEX2 proximity labeling for cell-type and subcellular compartment specific proteomics in the mouse brain. APEX is an engineered peroxidase that can be rapidly induced to tag proteins in minutes with biotin phenol (BP) and H_2_O_2_^12^. To broadly target major subcellular proteomes in the mouse brain, we created and expressed three Cre-dependent APEX variants (**Fig. 1a**) that localized to the nucleus, to the cytosol, and to the membrane compartments. Targeting was attained using well validated sorting motifs (Histone 2B fusion, H2B; nuclear exporting sequence, NES; and membrane anchor LCK sequence, respectively). We demonstrated Cre-dependent expression by neonatal viral transduction in the striatum of Drd1^Cre^ mice with AAV encoding APEX variants and a P2A-linked EGFP reporter (**Fig. 1a**). We confirmed Cre- dependent expression by examining the projection pattern of the EGFP reporter. Drd1^Cre^+ neurons project primarily to the SNr, as expected for this cell type^18^, in all APEX variants (**Fig. 1b and Supplementary Fig. 1a**). GFP expression was similar across constructs, as expected for P2A-linked EGFP, whose distribution is independent from the APEX localization and should be the same regardless of APEX targeting.

**Figure 1.**
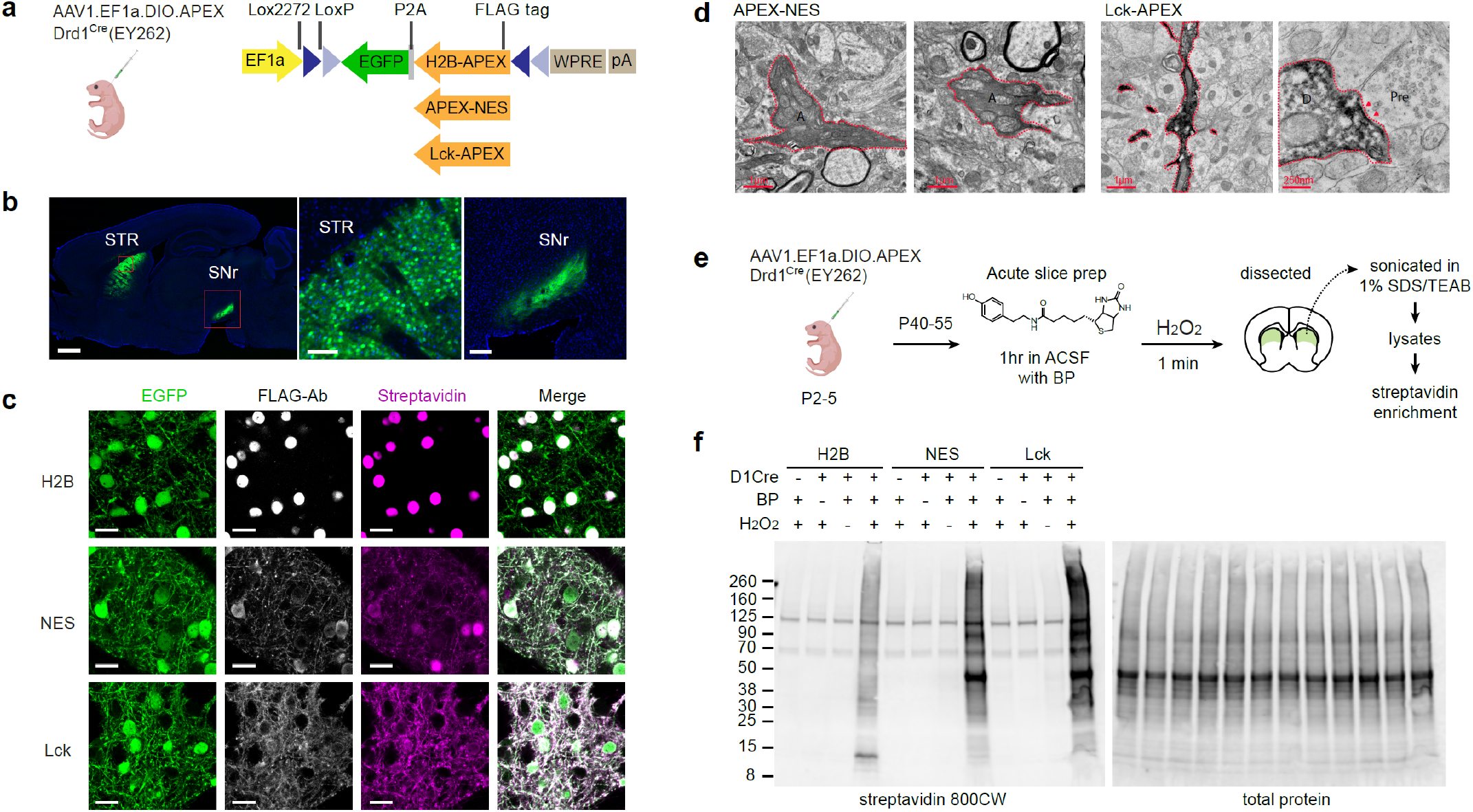
Genetically targeted subcellular protein labeling in the mouse brain. (a) Design of APEX constructs. *Left*, neonatal AAV transduction to selectively express APEX in Drd1^Cre^-positive striatal neurons. *Right*, FLAG-tagged APEX constructs targeting the nucleus (H2B), the cytoplasm (NES), and the membrane (LCK). All constructs contain a P2A-linked EGFP and are under EF1a promoter. Coding sequences were inverted and flanked by lox sites in a DIO-cassette for Cre-dependent expression. (b) Cre-dependent APEX-NES expression in the striatum of Drd1^Cre^ mouse line. *Left*, sagittal section (1mm) of Drd1^Cre^ > 2 weeks post AAV transduction. *Middle*, somatic-filled EGFP reporter was observed in the striatum (STR, 100 µm). *Right*, dSPN-specific axonal projection was observed in the SNr (200 µm). (c) Subcellular localization of APEX constructs in striatal neurons. Confocal images of APEX-expressing neurons including EGFP, immunostained FLAG-Ab, and streptavidin for visualizing APEX subcellular localization and biotinylation patterns (scale bar: 20 µm). (d) APEX labeling pattern under transmission electron microscopy. *Left*, APEX-NES containing axons (A) in the SNr show diffused labeling pattern across the cytosol. Right, Lck-APEX containing dendrite show dense membrane-enriched signal with visible intracellular organelles. A cross section of a dendritic spine (D) with a post-synaptic density (red arrow), juxtaposed to a pre-synaptic site (Pre). (e) *Ex vivo* biotinylation workflow for acute brain slices. APEX expression was achieved via neonatal AAV transduction. (f) Conditional protein biotinylation in acute brain slices. APEX-mediated biotinylation requires BP and H_2_O_2_.

To estimate APEX biotinylation pattern, we took advantage of retained APEX activity in paraformaldehyde (PFA)- fixed brain slices^19^. We performed protein biotinylation in PFA-fixed sections using biotin phenol. Biotinylated sections were immunostained for APEX and biotinylated proteins (**Fig. 1c)**. We created somata masks and computed the EGFP and streptavidin signal intensity inside and outside the masks. Untargeted EGFP has similar distribution among constructs, while biotinylated protein signal differs, revealing H2B with most signal inside cell bodies and LCK with most signal outside cell bodies (**Supplementary Fig. 1b**). In addition, a line scan across neuronal somata confirms that H2B-APEX labels proteins towards the nuclear compartment. APEX-NES broadly labels proteins in the somata, and LCK-APEX labels proteins away from the somata (**Supplementary Fig. 1c**). This is in an agreement with the absence of FLAG-Ab immunostained H2B signal in the SNr, supporting its restricted expression in nuclei (**Supplementary Fig. 1a**). To further distinguish the NES and LCK constructs, we utilized APEX-peroxidase activity in fixed brain slices to selectively deposit diaminobenzidine-osmium stain and inspect localization patterns under a transmission electron microscope^12^ (**Fig. 1d**). The NES construct showed a diffuse pattern filling axonal cross sections. In contrast, the labeling pattern for LCK in dendrites was more membrane enriched. Altogether, all APEX AAV constructs were Cre-dependent and localized to distinct subcellular compartments and were suitable for Cre-dependent ultrastructural analyses. Importantly, APEX construct expression did not alter basic electrophysiological properties in neurons expressing the constructs, relative to mCherry reporter controls (**Supplementary Fig. 2**).

APEX-based proteomics utilizes biotin phenol and hydrogen peroxide to label tyrosine residues via an oxidative process^20^. To ensure efficient delivery of biotin phenol in tissues, we developed an *ex vivo* biotinylation workflow optimized for acute brain slices (**Fig. 1e**). Drd1^Cre^ neonates were virally transduced with Cre-dependent APEX AAVs. Five to six weeks after transduction, 250 µm acute slices were prepared and incubated in carbogenated artificial cerebrospinal fluid (ACSF), supplemented with 500 µM biotin phenol for 1hr. Biotinylation was induced by transferring slices to ACSF containing 0.03% H_2_O_2_ for 1 min, followed by immersing slices in a quenching solution. EGFP-positive region in the dorsal striatum was dissected for western blot analysis (**Fig. 1f**). Western blot of total striatal lysates indicated that proteins were rapidly labeled within the 1 min H_2_O_2_ exposure time window. Biotinylated proteins in control samples were mostly accounted for by the endogenously biotinylated set. The pattern of H2B sample was distinguishable from that of NES and LCK (the presence and the absence of ∼15kD and ∼50kD bands, respectively), in an agreement with histology data.

To demonstrate the application of Cre-dependent APEX AAV strategy for cell-type and subcellular compartment specific proteomics, streptavidin magnetic beads were used to enrich biotinylated proteins from striatal lysates prepared separately for each animal (with no pooling) (**Fig. 2a**). Western blot analysis of input, flowthrough, and bead eluate confirmed a successful enrichment protocol (**Fig. 2b-c**). Streptavidin signal in lysate input was depleted in the flowthrough fraction, and the presence of signal in the bead eluate verified that biotinylated proteins were properly captured. Next, we digested protein-captured beads with trypsin to generate peptides for the bottom-up proteomics approach, where peptides are analyzed by liquid-chromatography-coupled tandem mass spectrometry (LC MS/MS). To generate MS data, peptides were labeled with 10plex TMT reagents in a complete randomized block design (**Fig. 2a, Supplementary Fig. 3a)**. The total of 20 samples were randomized across two 10-plex sets (four biological replicates for Cre-negative control, H2B+, NES+, and LCK+, with reference-normalization samples for each 10-plex).

**Figure 2.**
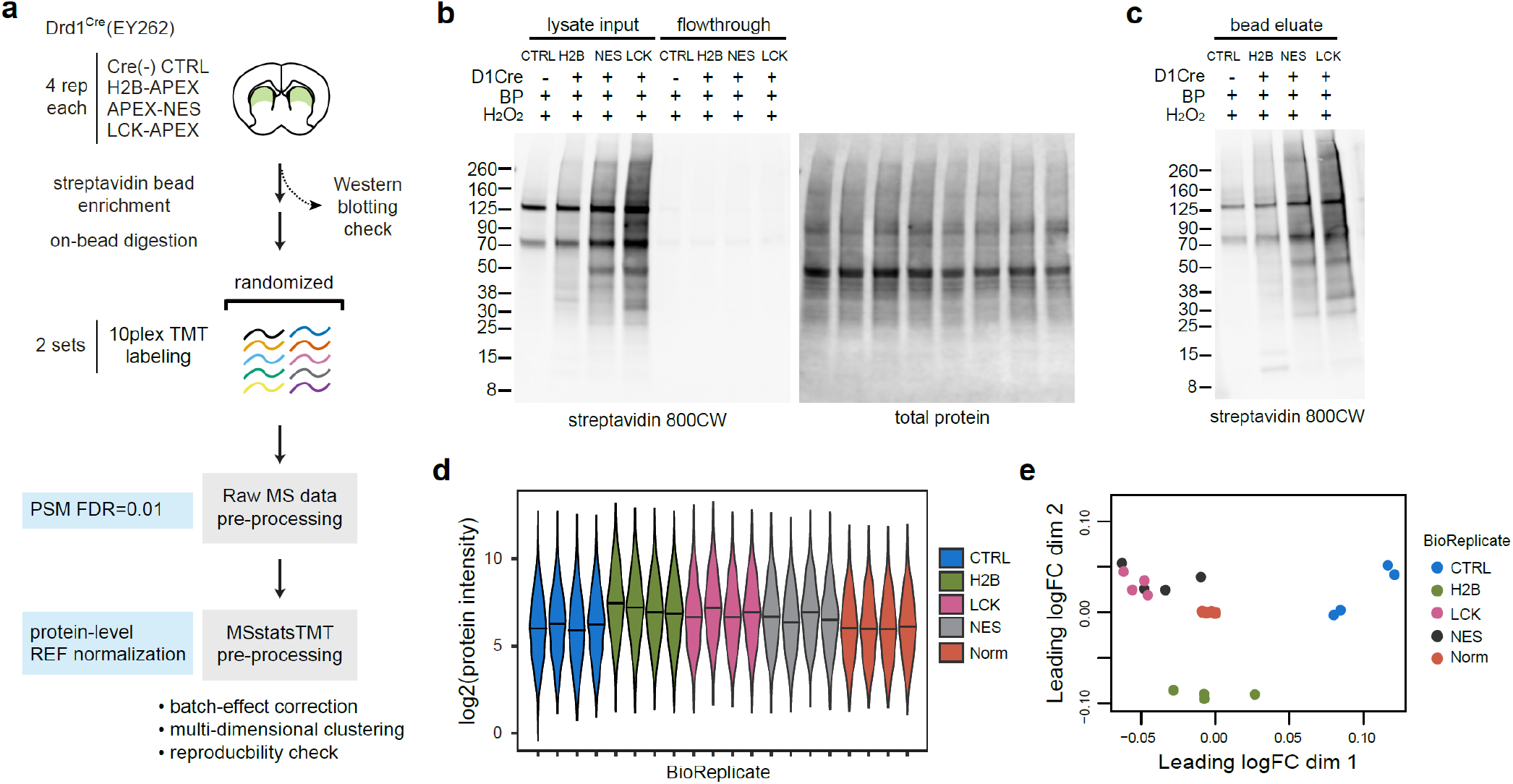
APEX2-based cell-type specific proteomics in the mouse striatum. (a) Workflow for proteomics sample preparation. Streptavidin bead enrichment following brain lysate preparation was used to isolate biotinylated proteins for MS analysis. (b) Western blot analysis of biotinylated brain lysate and flowthrough fractions. Equal amount of proteins and streptavidin beads are used in the enrichment process. Depletion of biotinylated in the flowthrough fractions confirms successful enrichment. (c) Western blot analysis of eluate fraction after streptavidin bead enrichment. Differential enrichment output reflects varying amount of biotinylated proteins from each sample. (d) Overall protein intensities across biological replicates. (e) Multidimensional scaling plot approximates expression differences between the samples for the top 500 proteins.

In this experimental setup, peptide sequences were identified using the MS2 spectra, while quantification of reporter ions in each TMT channel was performed using the SPS MS3 method with real-time search^21^. Peptide spectral matches (PSMs) were done in Proteome Discover by SEQUEST search engine with 1% false discovery rate (FDR). We identified a total of 1552 proteins in across both 10-plex experiments. We used MSstatsTMT R package to perform protein-level summarization and quantification^22^. A total of 736 proteins with unique peptides were quantified. The overall log2 protein intensities for each biological replicate were plotted in **Fig. 2d**. TMT batch effects were corrected protein-by-protein based on protein signal in the reference channel. A multidimensional scaling (MDS) plot after reference-channel normalization revealed that APEX construct replicates cluster together (**Fig. 2e**). This normalization is necessary in TMT experiments: an MDS plot of un- normalized data showed a strong separation between two TMT batches (**Supplementary Fig. 3b**). To evaluate the reproducibility of this workflow, we plotted an example of a multi-scatter of H2B biological replicates. Pearson correlation of log2 protein intensities were highly correlated (r > 0.98) (**Supplementary Fig. 3c**). The distribution of coefficient of variation percentages (%CV) for each condition were similar, with a mean(%CV) of < 10% (**Supplementary Fig. 3d-e**). Collectively, biotinylation, streptavidin enrichment, and MS sample preparation process was highly reproducible.

Genetically targeted proximity labeling approaches offer information on cell-type specificity and subcellular localization. To determine whether identified proteins were cell-type specific, we defined the first filter based on comparisons between Cre-positive samples and Cre-negative control samples using a pairwise moderated t-test implemented in MSstatsTMT (**Fig. 3a**). Proteins that were not statistically significantly enriched in the positive direction were discarded as non-specific binders (**Fig. 3b**). Some examples included known endogenously biotinylated proteins (Acaca, Pccb, Pcca, Pc, Mccc1). Retained proteins with log2-fold change > 0 and adjusted p-values < 0.05 were considered as cell-type specific and enriched due to the biotinylation process. In an agreement with western blot and histology, the majority of proteins detected in Cre-positive samples were retained as cell-type specific (**Fig. 3b, Supplementary Fig. 4a-b**). Next, we aimed to define a second, stringent cutoff for the nuclear and membrane-enriched proteome using t-tests (**Fig. 3c**). We performed a pairwise comparison for H2B-NES and H2B-LCK to generate nuclear-enriched and membrane-enriched protein list, respectively. We obtained 74 nuclear enriched proteins and 26 membrane-enriched proteins after applying the second filter. The choice of a reference compartment was important for the statistical comparison. NES-LCK comparison did not yield any statistically significant enrichment for either compartments (adj. p > 0.05). This was mostly because proteins labeled by the NES and LCK were accessible to biotin phenol, with a majority of proteins localizing to both membrane and cytoplasm (**Supplementary Fig. 4c**). A rank plot between NES and LCK indicated a relative enrichment of membrane proteins towards the LCK construct. This suggests that our method is fairly conservative. Alternatively, a rank-based or static cutoff can be used instead^23^ (**Supplementary Fig. 4c- d**).

**Figure 3.**
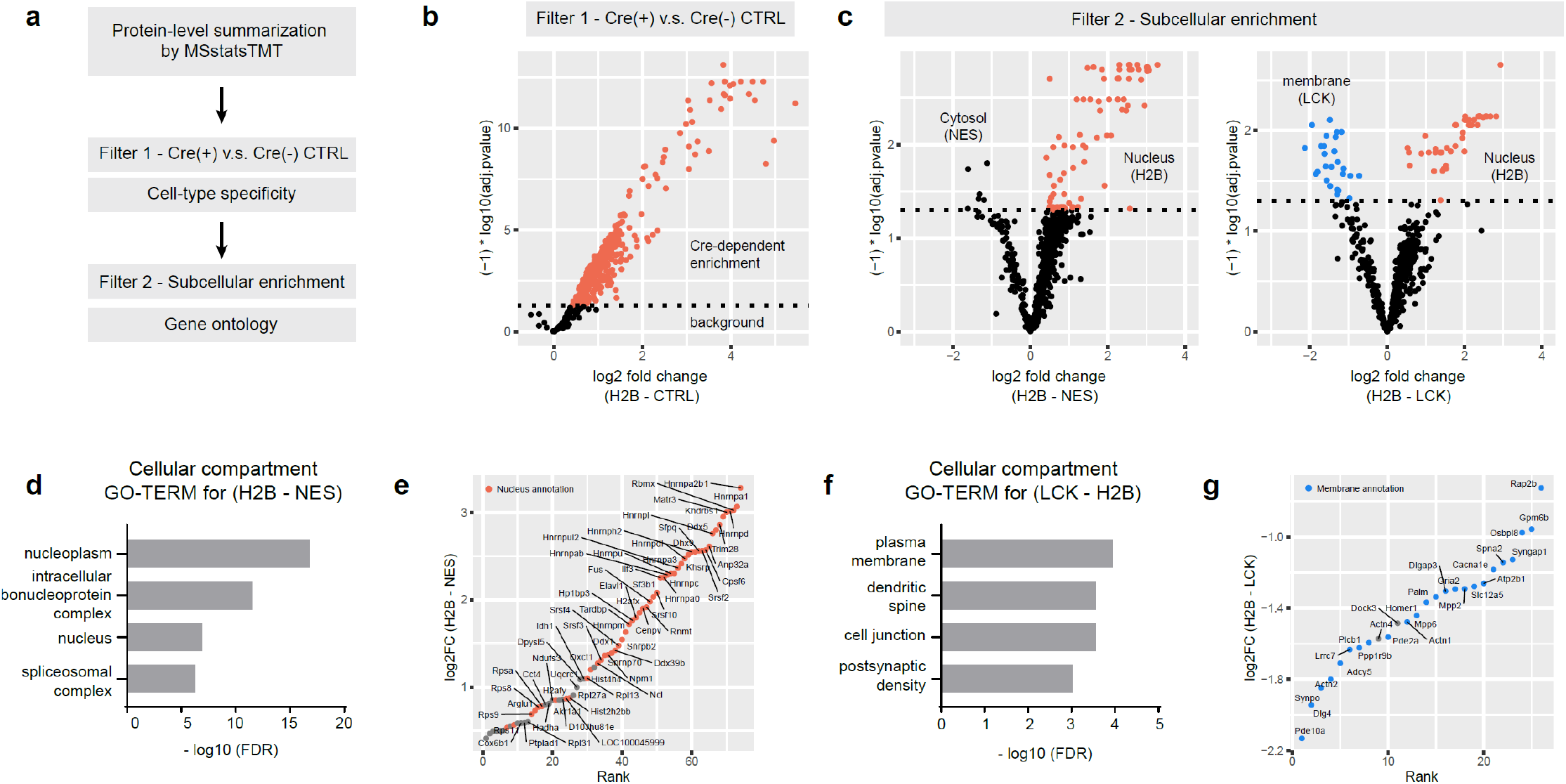
*In situ* proximity labeling generates cell-type and subcellular compartment specific proteomes. (a) Statistical comparison strategy. Moderated t-test for significance with Benjamini-Hochberg (BH) multiple hypothesis correction was used as a cut-off. Filter 1 was the first comparison to determine the degree of protein enrichment above non-specific background. Cre-positive samples were compared against Cre-negative control. Filter 2 was used with proteins that were retained after the first filter to determine the extent of subcellular enrichment. (b) Volcano plot for H2B v.s. CTRL. Proteins that were significantly enriched above background were indicated by red dots (adj. p < 0.05). Proteins below the dotted line of adj. p = 0.05 were considered background. (c) Volcano plots for H2B v.s. NES and H2B v.s. LCK. *Left*, nuclear enriched proteins (H2B) were indicated by red dots. *Right*, nuclear enriched proteins (H2B), red dots; membrane-enriched proteins (LCK), blue dots. Proteins below the dotted line of adj. p = 0.05 were differentially enriched but not statistically significant. (d) Gene ontology cellular component analysis for H2B-enriched proteins. (e) Log2FC-rank plot for H2B-enriched proteins. Proteins with UNIPROT nucleus annotation, red dots. Positive log2 fold change (H2B-NES) indicates biased enrichment towards the nucleus. (f) Gene ontology cellular component analysis for LCK-enriched proteins. (g) Log2FC-rank plot for LCK-enriched proteins. Proteins with UNIPROT membrane annotation, blue dots. Negative log2 fold change (H2B-LCK) indicates biased enrichment towards the membrane.

To examine whether H2B-enriched and LCK-enriched proteins are overrepresented by any particular subcellular compartments, we used gene ontology analysis with total identified proteins as background. Indeed, the top four terms with lowest FDR for each protein list were nucleus- and membrane-related terms, confirming that the two APEX constructs labeled proteins in different compartments (**Fig. 3d, 3f**). Next, we created log2 foldchange vs rank plots with nucleus- or membrane annotations. As expected, proteins with greater fold change are more likely to have prior UNIPROT nucleus or membrane annotations for H2B and LCK, respectively (**Fig. 3e, 3g**). Consistent with recent proteomic profiling of Drd1+ striatal nuclear proteome using fluorescent nuclei sorting, we found that Drd1+ nuclear proteome preferentially expressed Hnrnpa2b1 and Hnrnpd protein network^4^. In the H2B rank plot, a few proteins (*e*.*g*., Oxtc1, Idh1, Ndufs3) showed mitochondrion annotations. This was not surprising because mitochondrial compartment is oxidative. Thus, it is likely that an APEX-based approach could enrich mitochondrial proteins in addition to proteins from the intended compartments. In the LCK rank plot, there were two proteins (Dock3 and Actn4) that do not list membrane as their primary cellular component annotation in the UNIPROT database. Dock3 is a member of guanine nucleotide exchange factors (GEFs) that regulates membrane-associated protein, Rac1. It was also shown to interact with NR2B subunit of NMDA receptors^24^. Actn4 is a filamentous actin-binding protein containing a group1-mGluR binding domain^25^. It has been shown that Actn4-mGluR interaction was involved in remodeling of dendritic spine morphology^25^. Although Dock3 and Actn4 distribute throughout the cytoplasm, both were known to interact with membrane proteins, suggesting a potential application of this workflow for *in situ* protein-protein interaction studies.

In this study, we demonstrated a cell-type specific APEX2 proximity labeling workflow to profile the subcellular proteome in the mouse brain across the nucleus, the cytoplasm, and the intracellular membrane compartment. As proof of principle, we focused on the Drd1^Cre^-positive neurons in the striatum, because the data generated here could be used as a reference for comparison with publicly available scRNAseq^2^, TRAPseq^26^, and nuclear proteome^4^ data of the same cell type. From a technical standpoint, our Cre-dependent AAV transduction can be applied to many available Cre-driver mouse lines in a neural circuit of interest. Although we used unpooled striatal lysates from one brain as an input in the streptavidin enrichment process, pooling across animals may be necessary for smaller brain regions or very low abundance subcellular compartments. In addition, peptide fractionation can be used to increase the depth of identification and quantification.

The APEX workflow offers subcellular specificity and speed. It has been shown that there are distinctions among cell-type organellar proteomes in the brain^27^. This raises a question about the homeostatic capacity across neuronal types in response to environments. APEX-based proteomics is a good candidate for mapping the proteome of cell-type organelles including compartments that cannot be isolated with conventional biochemical fractionation methods^17,23,28^. In addition, APEX labeling reaction in the brain slices is efficient, with a 1 min time window. Although delivery of biotin phenol requires an *ex vivo* incubation, APEX labeling speed provides a spatiotemporally resolved method to study genetically targeted proteome, also unlocking access to transient protein-protein interaction information^29,30^. The expansion in the availability of Cre driver lines for rodents and growing genetic traction over previously inaccessible animal models, along with the increase in the number of APEX toolkits^3,19^ and the new validated workflow presented here, positions APEX-based proteomics to become the major cell type-specific proteomics approach for diverse neuroscience applications.

## Methods

### Plasmid construction and AAV preparation

APEX2-NES-P2A-EGFP, LCK-APEX2-P2A-EGFP, and H2B-APEX2-P2A-EGFP were synthesized by Genscript and subcloned into pAAV-EF1a-DIO-WPRE-hGH vector (a gift from Dr. Karl Deisseroth (Addgene plasmid #20297) using restriction enzymes AscI and NheI. APEX2 sequence was cloned based on pcDNA3-APEX2- NES (a gift from Dr. Alice Ting; Addgene plasmid #49386^12^). APEX AAVs was packaged into adeno-associated virus serotype 1 by the University of North Carolina (UNC) vector core service or Vigene Biosciences (Rockville, MD. USA). AAV8.hSyn.DIO.Cherry was a gift from Dr. Bryan Roth (Cat. No. 50459-AAV8, Addgene).

### Mouse strains and genotyping

Animals were handled according to protocols approved by the Northwestern University Animal Care and Use Committee. Weanling and young adult male and female mice were used in this study. Drd1^Cre^ (262Gsat/Mmcd)^31^ was obtained from Mutant Mouse Regional Resource Center (MMRRC) at the University of California, Davis. C57BL/6 mice used for breeding and backcrossing were acquired from Charles River (Wilmington, MA). All mice were group-housed, with standard feeding, light-dark cycle, and enrichment procedures; littermates were randomly assigned to conditions. All animals were genotyped according to the MMRRC strain-specific primers and protocols using GoTaq Green PCR master mix (Cat. No. M712, Promega Corporation, Madison, WI, USA).

### Stereotactic injections

Conditional expression of APEX and reporters in Cre+ neurons was achieved by recombinant adeno-associated viral transduction encoding a double-floxed inverted open reading frame (DIO) of target genes, as described previously. For neonatal AAV delivery, P3-6 mice were cryoanesthetized and were placed on a cooling pad. For all APEX constructs, 400 nl of AAV were delivered using an UltraMicroPump (World Precision Instruments, Sarasota, FL). Dorsal striatum was targeted in neonates by directing the needle +0.1 mm anterior of bregma, ±0.3 mm from midline, and 1.8-2.0 mm ventral to skin surface. Following the procedure, pups were warmed on a heating pad and returned to home cages. AAVs were diluted to the final titers using Gibco PBS pH 7.4 (AAV1.EF1a.DIO.APEX-NES-P2A-EGFP, AAV1.EF1a.DIO.LCK-APEX-P2A-EGFP, AAV1.EF1a.DIO.H2B- APEX-P2A-EGFP titer ∼ 3⨯10^12^ GC/ml). P35-P70 animals were used for histology, western blots, and proteomics experiments.

### Acute slice preparation and electrophysiology

Acute slice preparation was adapted from a previously published protocol^32^. Animals were anesthetized with isoflurane and perfused with ice-cold artificial cerebrospinal fluid (ACSF) containing (in mM): 127 NaCl, 25 NaHCO_3_, 1.25 H_2_Na_2_PO_4_ monobasic, 25 D-Glucose, 2.5 KCl, 1 MgCl_2_, and 2 CaCl_2_ (osmolarity ∼310 mOsm/L). Animals were decapitated, and the brain was immediately removed and submerged in ice-cold ACSF. Tissue was blocked using a 4% agar block and transferred into a slice chamber containing ice-cold ACSF bubbled with 95%O_2_/5%CO_2_. Bilateral coronal slices (300 μm) were cut on a Leica VT1000 S slicer. Slices were cut in lateral-medial direction and transferred into a holding chamber containing pre-warmed (34°C) and oxygenated. Slices were incubated at 34°C for 15 minutes and recovered at RT for 30 minutes.

Slices were transferred into a recording chamber perfused with oxygenated ACSF at a flowrate of 2-4 mL/min at RT. Whole-cell patch-clamp recordings were conducted on spiny projection neurons (SPNs). Patch pipettes with ∼5-8 MΩ resistance were filled with internal solution containing (in mM): 115 K-Gluconate, 20 KCl, 4 MgCl_2_, 10 HEPES, 4 Mg-ATP, 0.3 Na-GTP, 7 Phosphocreatine (disodium salt hydrate), and 0.1 EGTA (in KOH) (pH 7.2, osmolarity 290 mOsm). Excitability experiments were conducted in current clamp mode, where holding current was set to hold cells at approximately -70mV. Recordings were made using a 700B amplifier (Axon Instruments, Union City, CA); data were sampled at 10 kHz and filtered at 4 kHz with a MATLAB-based acquisition script (MathWorks, Natick, MA). Series and input resistance were monitored using a 200 ms, -5 pA pulse at the end of every sweep. Acquisition intervals were 20 sec long; sweeps were 5 sec long with 250 ms long current injections after a 200 ms long delay. 200 pA current injection amplitude was injected (4 sweeps/cell minimum).

### Quantification and statistical analysis for electrophysiology data

Offline analysis of electrophysiology was performed using IgorPro (Wavemetrics, Portland, OR). Action potential shape analysis was performed using a MATLAB-based analysis script. Sex and age were balanced across groups. Statistical analysis was performed using GraphPad Prism software (GraphPad, LaJolla, CA). Group data are expressed as group means ±SEM. Multiple group comparisons were done using one-way ANOVA with Bonferroni correction. Adjusted p < 0.05 was considered statistically significant.

### Tissue processing and immunohistochemistry

Mice were deeply anaesthetized with isoflurane and transcardially perfused with 4% paraformaldehyde (PFA) in 0.1 M phosphate buffered saline (PBS). Brains were post-fixed for 1-5 days and washed in PBS, prior to sectioning at 50-80 µm on a vibratome (Leica Biosystems). To verify APEX activity and subcellular localization, tissue sections were incubated in PBS containing 500 µM biotin phenol for 30 min and treated with PBS containing 0.03% H_2_O_2_ for 1min. The reaction was quenched 3x with PBS containing 10 mM NaN_3_ and 10mM sodium ascorbate. For other immunostaining, no biotinylation was performed. Sections were incubated with primary antibody with 0.2% Triton X-100 for 24-48 hrs at 4°C. On the following day, tissues were rinsed three times with PBS, reacted with secondary antibody for 2 hrs at RT, rinsed again for three times. Sections were mounted dried on Superfrost Plus slides (Thermo Fisher, Waltham, MA), air dried, and cover slipped under glycerol:TBS (9:1) with Hoechst 33342 (2.5 µg/ml, Thermo Fisher Scientific). Primary antibodies used in the study were mouse anti-FLAG (1:1000; Cat. No. A00187-200, Genscript, NJ, USA), chicken anti-GFP (1:2000; Cat. No. AB13970, Abcam, Cambridge, UK). Alexa Fluor 647-conjugated secondary antibodies against mouse, chicken and/or Alexa Fluor 647-conjugated streptavidin (Life Technologies, Carlsbad, CA) were diluted to 1:500 for all secondary antibody staining steps. Whole sections were imaged with an Olympus VS120 slide scanning microscope (Olympus Scientific Solutions Americas, Waltham, MA). Confocal images were acquired with a Leica SP5 confocal microscope (Leica Microsystems).

FIJI was used for APEX labeling quantification. For calculating the ratio of fluorescent signal inside v.s. outside the cell body, somata masks were manually drawn for each confocal image frame in the GFP channel. The same mask was applied to streptavidin channel. For vertical line scan analysis, random cells across multiple confocal images were selected, fluorescent intensity for GFP and streptavidin were normalized to the corresponding maximum intensity within a given line scan and averaged across 10 cells. Statistical analysis was performed using GraphPad Prism software (GraphPad, LaJolla, CA). Group data are expressed as group means ±SEM. Multiple group comparisons were done using one-way ANOVAs with Tukey’s correction. Adjusted p < 0.05 was considered statistically significant.

### Transmission electron microscopy (TEM)

Transcardial perfusion and biotinylation were performed as described above except for the fixatives. For TEM specimen preparation, 2% glutaraldehyde and 2% PFA in PBS was used. 100 µm brain slices were washed several times with 0.05 M sodium phosphate buffer (PB), and then processed for TEM with 2 exchanges of the primary fixative that consisted of 2.5% glutaraldehyde, 2% PFA in 0.1 M PB. The brain slices were again washed 3x with the buffer followed by a secondary fixation in 1.5% osmium tetroxide (aqueous). Samples were washed 3x with DI water before beginning an acetone dehydration series. All of the preceding steps up were carried out in Pelco Biowave Microwave with Cold Spot and vacuum. EMBed 812 embedding media by EMS was gradually infiltrated with acetone for flat embedding. The selected ROI was cut out and mounted on a blank stub for sectioning. 90 nm thin sections were collected on copper grids using a Leica Ultracut S ultramicrotome and DiATOME 45° diamond knife. Images were acquired at 100kV on a 1230 JEOL TEM and Gatan Orius camera with Digital Micrograph software.

### *Ex vivo* biotinylation for proteomics studies

Acute slices were prepared as described above. Viral expression in 250 µm-thick striatal slices were confirmed by NightSea Dual FP flashlight. For *ex vivo* biotinylation, slices were incubated in carbogenated ACSF with 500 µM biotin phenol (Iris Biotech) at RT for 1 hour. They were briefly rinsed in ACSF and then transferred into ACSF containing 0.03% H_2_O_2_ for 1min. The reaction was quenched by transferring slices to ASCF containing 10 mM NaN_3_ and 10 mM sodium ascorbate. Fluorescent-positive regions were dissected in ice-cold ACSF under a fluorescent dissection microscope (NIGHTSEA, Lexington, MA, USA) and transferred to 1.5 ml polypropylene tubes. Tissues were frozen and stored at -80°C for further processing.

### Protein extraction, streptavidin enrichment, and on-bead trypsin digestion

Total protein was extracted by sonication in 400 µl lysis buffer (1% SDS, 125mM TEAB, 75mM NaCl, and Halt™ protease and phosphatase inhibitors). Lysates were cleared by centrifugation at 12000 g for 15min at 4°C. Supernatant was used for subsequent procedure. Total protein (1 µl of samples were diluted to 100 µl with water) was estimated using microBCA assay according to the manufacturer instructions (Cat. No. 23235, Thermo Fisher). 300 µg of brain lysates (in 200 µl) were reduced with 20 µl 200 mM DTT for 1 hr and alkylated with 60 µl 200 mM IAA in the dark for 45 min at 37°C with shaking. 240 µl of reduced/alkylated lysates were incubated with 300 µl of streptavidin magnetic beads (prewashed twice with 1 mL no-SDS lysis buffer for 1 hr at RT with shaking. Enriched beads were washed with 2x 1 mL no-SDS lysis buffer, 1 mL 1M KCl, 5x 1 mL no-SDS lysis buffer (5 min each). Washed beads were digested with trypsin/LysC solution (∼6 µg in 100 µl 100 mM TEAB) overnight at 37°C. Digested supernatant was collected. Beads were rinsed with 50 µl 100 mM TEAB. Supernatant were combined. Trace amount of magnetic beads were removed twice by magnetization and tube changes. 10 µl from each sample was saved for Pierce fluoremetric peptide quantification assay (Cat. No. 23290, Thermo Fisher), and 16 µl from each sample were combined to make an average reference sample for TMT batch normalization. Peptides were frozen and dried in a vacuum concentrator before TMT labeling.

### TMT-labeling, Fractionation, and Desalting

Dried peptide samples were reconstituted in 20 µl 100 mM TEAB and sonicated for 15 min at RT. TMT labeling protocol was performed similar to Zecha et al. (20 µl peptides + 5 µl 59 mM TMTzero)^33^. Briefly, one set of 0.8 mg 10plex TMT reagents was warmed up to room temperature and dissolved in 41 µl Optima LC/MS-grade acetonitrile (ACN). 5 µl of ∼56 mM TMT reagents was added to 20 µl reconstituted peptide according to the experimental design in **Supplemental Fig 2b**. Labeling was performed at RT for 1 hr with shaking at 400 rpm. The reaction was quenched by adding 2.2 µl 5% hydroxylamine/100 mM TEAB at RT for 15 min with shaking. Samples were mixed equally, acidified to pH < 3 with formic acid (FA) and dried in a speed vacuum concentrator. TMT mixture was resuspended in 50 µl buffer A (LCMS water 0.1% FA) and desalted using Pierce C18 spin tip (Cat. No. 84850, Thermo Fisher). All centrifugation steps were performed at 1000 g for 1 min. Spin tips were activated twice using 20 µl 80% ACN 0.1% FA and equilibrated twice using 20 µl buffer A. Samples were loaded 10 times. Spin tips were washed twice with 20 µl buffer A. Peptides were eluted with 40 µl 80% ACN 0.1% FA and dried in a vacuum concentrator and stored at -80°C until MS data collection.

### Mass spectrometry data acquisition and raw data processing

∼1 ug of TMT labeled peptides were resuspended in 2% acetonitrile/0.1% formic acid prior to being loaded onto a PepMap RSLC C18 2 µm, 100 angstrom, 75µm x 50 cm column (ThermoScientific) and eluted over a 90- minute gradient from 98% buffer A (H_2_O, 0.1% formic acid) to 45% buffer B (Acetonitrile, 0.1% formic acid). Sample eluate was electrosprayed (2000V) into a Thermo Scientific Orbitrap Eclipse mass spectrometer for analysis. The instrument was operated in MS3 with real-time search (max search time = 34 s; Xcorr = 0.1; ppm = 5) and synchronous precursor selection. MS1 spectra were acquired at a resolving power of 120,000. MS3 spectra are acquired at a resolving power of 60,000 with a max injection time of 400 ms.

Raw MS files were processed in Proteome Discoverer version 2.4 (Thermo Scientific, Waltham, MA). MS spectra were searched against the *Mus musculus* SwissProt database. SEQUEST search engine was used (enzyme=trypsin, max. missed cleavage = 4, min. peptide length = 6, precursor tolerance=10ppm). Static modifications include acetylation (N-term, +42.011 Da), Met-loss (N-term, -131.040 Da), Met-loss+Acetyl (N- temr, -89.030 Da), and TMT labeling (N-term and K, +229.163 Da). Dynamic modifications include oxidation (M, +15.995 Da) and biotinylation (Y, +361.146 Da). PSMs were filtered by the Percolator node (max Delta Cn = 0.05, target FDR (strict) = 0.01, and target FDR (relaxed) = 0.05). Reporter ion quantification was based on corrected S/N values with the following settings: integration tolerance = 20ppm, method=most confident centroid, co-isolation threshold = 70, and SPS mass matches = 65. PSMs results from Proteome Discoverer were exported for analysis in MSstatsTMT R package (version 1.7.3).

### Western Blotting

Tissue lysis and biotinylated protein enrichment procedure were described in method section above. Protein lysate inputs or flowthrough fractions were mixed with 6x laemmli loading buffer and heated to 90-95°C for 10 min. For eluting biotinylated protein off the beads, washed beads were mixed with 20 µl of 2x laemmli buffer containing 25 mM TEAB, 75 mM NaCl, and 20 mM biotin. Beads were heated to 90-95°C for 10 min. Proteins were separated in 4-20% gradient gels (Cat. No. 4561096, Biorad, CA, USA) and transferred to nitrocellulose membrane (Cat. No. 926-31090, LI-COR, NE, USA). Blots were briefly rinsed with TBS. For detection of biotinylated proteins, blots were incubated in TBST (0.1% tween-20) containing streptavidin CW800 (1:10000, Cat. No. 926-32230, LI-COR) for 1 hr at RT. Blots were washed three times with TBST for 10 min each. Total protein was detected using REVERT 700 according to the manufacturer instructions. Blots were scanned using a LI-COR Odyssey CLx scanner. All quantification was performed using LI-COR Image Studio version 5.2.

### Statistical analysis with MSstatsTMT, and gene ontology analysis

Peptide-level output including medium- and high-confidence peptides was exported from Proteome Discoverer and converted into MSstatsTMT-compatible format using PDtoMSstatsTMTFormat function. Only unique peptides were used for protein summarization with the following arguments: method = LogSum, global median normalization = FALSE, reference normalization = TRUE, imputation = FALSE. To reflect enrichment differences across the APEX constructs at the protein level, global median normalization was not performed. It was necessary to perform reference sample normalization to correct for TMT batch effects^34^. MSstatsTMT implements this correction protein-by-protein.

Multidimensional scaling was done using the plotMDS function in edgeR version 3.30.3^35^ using the default setting (top 500 proteins). Entries with missing values were omitted in this clustering. Coefficient of variations were calculated on log2 protein intensities. Multiscatter plots were generated using pairs.panels function in psych version 2.0.7. The mean values of log2 protein intensity were used to generate the log2 ratio histograms and scatter plots between Cre-positive samples (H2B, NES, LCK) and Cre-negative control.

Pairwise differential expression analysis was performed using moderated t-tests with Benjamini-Hochberg (BH) multiple hypothesis correction. Adjusted p-values < 0.05 were considered as statistically significant.

For gene ontology analysis, DAVID v6.8^36^ (https://david.ncifcrf.gov/) was used with identifier = ‘UNIPROT_ACCESSION’. All identified proteins in the dataset were used as background. Proteins with BH adjusted p-values < 0.05 and log2FC > 0 were used as gene list. Log10(FDR) was plotted for top four GO- TERMs.

## Data availability

Raw data that support the findings of this study are included in Supplementary Materials or will be made available upon request. Raw MS data will be deposited in the PRIDE database upon publication.

## Code availability

Analysis code will be made available upon request or available on Github.

## Acknowledgements

The authors are grateful to Lindsey Butler for mouse colony management, Northwestern Biological Imaging Facility and Dr. Tiffany Schmidt for confocal microscope access, Northwestern University BioCryo Facility for TEM sample preparation and microscope, Northwestern High throughput analysis laboratory for a microplate reader.

This work was supported by the NIMH R56MH113923, NSF CAREER Award 1846234, NINDS R01NS107539, the Beckman Young Investigator Award, Searle Scholar Award, Rita Allen Foundation Scholar Award, and Sloan Research Fellowship (all Y.K.), and NIMH R01MH118497 (M.L.M.). V.D. is a predoctoral fellow of the American Heart Association (19PRE34380056) and as an affiliate fellow of the NIH 2T32GM15538.

## Author contributions

V.D. and Y.K. designed the study. V.D. carried out most experiments and analyses in the study. V.D. created new plasmids, viruses, MS sample preparation and contributed to experimental analyses. G.S. made major contributions to electrophysiology experiments and analyses. M.L.M. made major contributions to MS data acquisition and analyses. V.D., M.L.M., and Y.K. wrote the manuscript.

## Supplementary information

Supplementary Text and Figures 1-4

