## Supplementary Figures 1-4 for "Cell type and subcellular compartment specific APEX2 proximity labeling proteomics in the mouse brain"

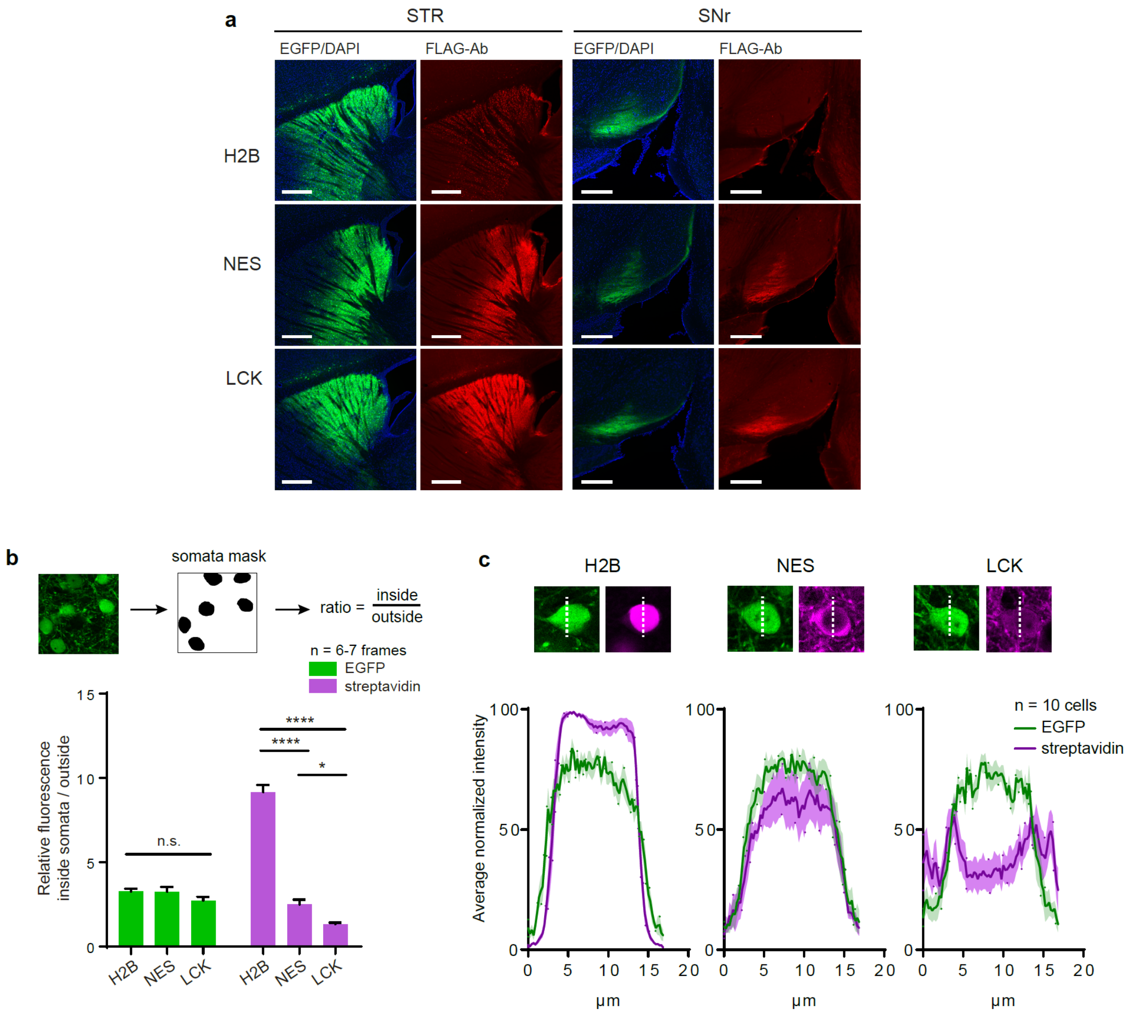


### Supplementary Figure 1. Subcellular targeting of the APEX constructs.

(a) Cre-dependent expression patterns of APEX variants in the striatum and the SNr. EGFP, DAPI, immunostained FLAG-Ab for APEX were in green, blue, and red, respectively (scale bars: 400µm)

(b) Distribution of APEX-dependent biotinylated proteins. *Top*, an example of somato masks used in ratio calculation for each image frame. *Bottom*, summary data for relative fluorescence signal inside and outside somata: green for EGFP, and magenta for streptavidin (n=6-7 per construct). One-way ANOVA with Tukey's multiple comparisons test: EGFP, $F(2,17) = 1.688$, $p = 0.2145$, streptavidin, $F(2, 17) = 185.3$, $p < 0.0001$. H2B vs NES, adj. $p < 0.0001$, H2B vs LCK, adj. $p < 0.0001$, and NES vs LCK, adj. $p = 0.0435$.

(c) Line scan analysis of biotinylated proteins across neuronal somata. *Top*, an example of a vertical line scan drawn across an APEX-expressing neuron. *Bottom*, fluorescent signal in the EGFP and streptavidin channels were plotted and normalized to the maximum intensity (n=10 cells/construct).

*p < 0.05, ** p < 0.01, *** p< 0.001. Error bars reflect SEM.

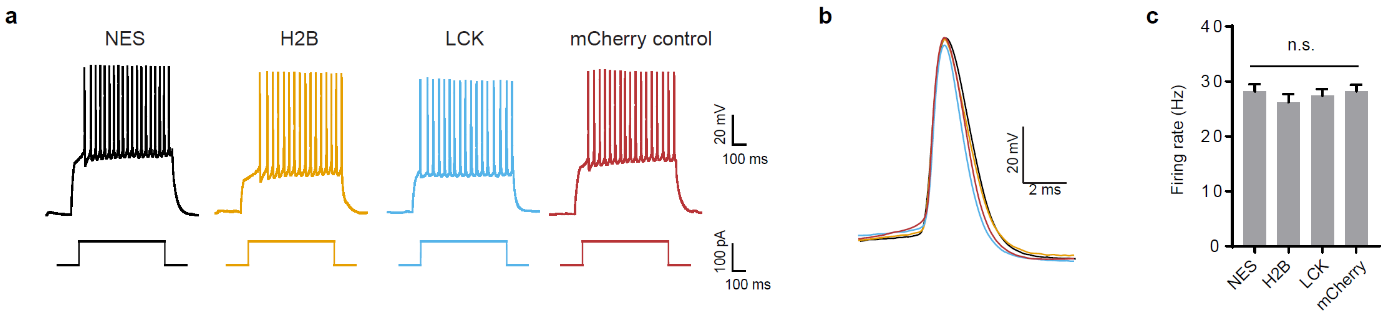


**Supplementary Figure 2. Electrophysiological characterization of APEX-expressing neurons.**

(a) Current-clamp recording of APEX-expressing dSPNs.
(b) Examples of action potential shape across APEX constructs.
(c) Summary for dSPN firing rate (Hz) with 200 pA current injections. One-way ANOVA with Bonferroni multiple comparisons test: F (3, 140) = 0.5109, p = 0.6754.

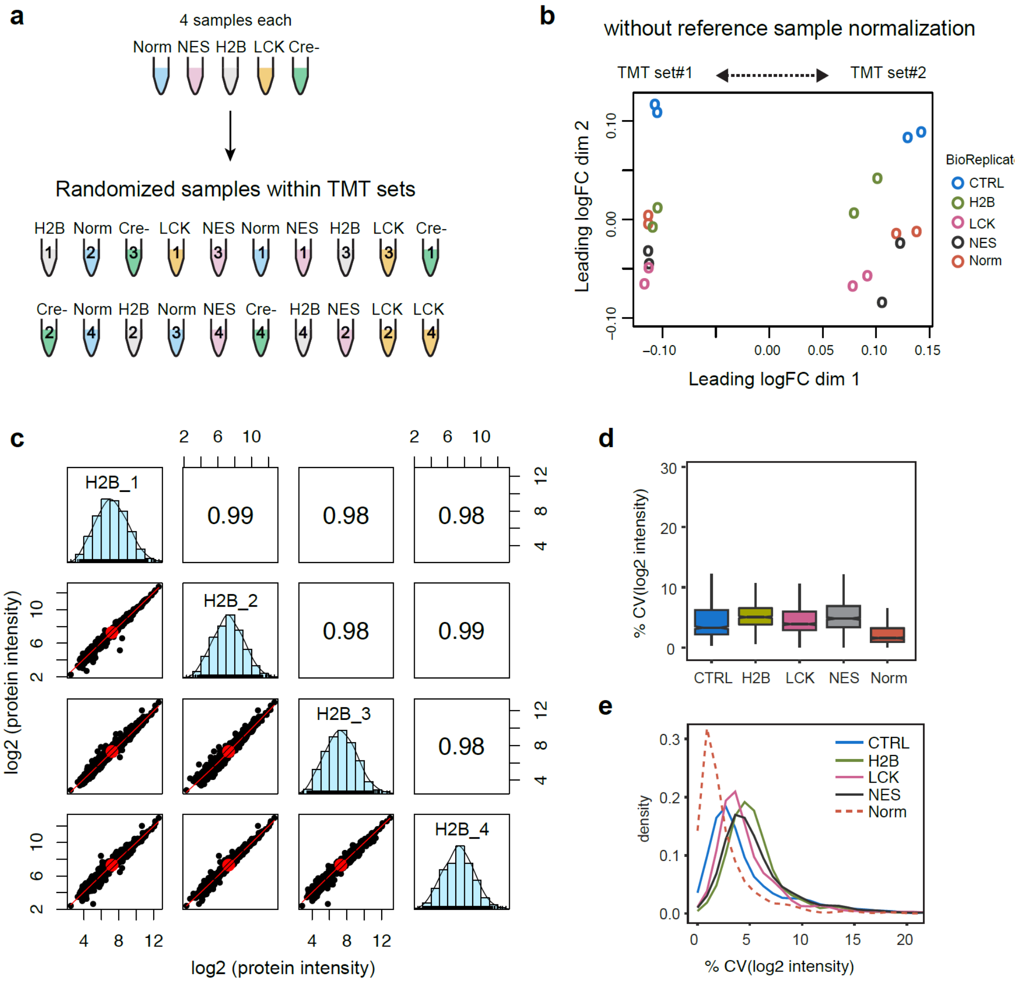


**Supplementary Figure 3. Analysis of MS sample preparation reproducibility.**

(a) Experimental design schematic. Samples were randomized across two 10plex TMT sets (top and bottom). Direction left-to-right was ordered by TMT channels (126-131). Each set contained 2 complete blocks (each block = Norm, NES, H2B, LCK, Cre-).

(b) Multidimensional clustering of log2 protein intensities without reference sample normalization. Each point was a biological replicate. Arrow indicates a clear separation between TMT sets, when reference normalization was not performed.

(c) Multiscatter plot of H2B biological replicates (example). *Diagonal*, histograms of protein log2 intensities. *Upper right corner*, Pearson correlation coefficient *r*, *Lower left corner*, pairwise scatter plots between replicates, red lines are loess-fit lines.

(d) Box plot of %CV distribution.

(e) Density plot of %CV distribution.

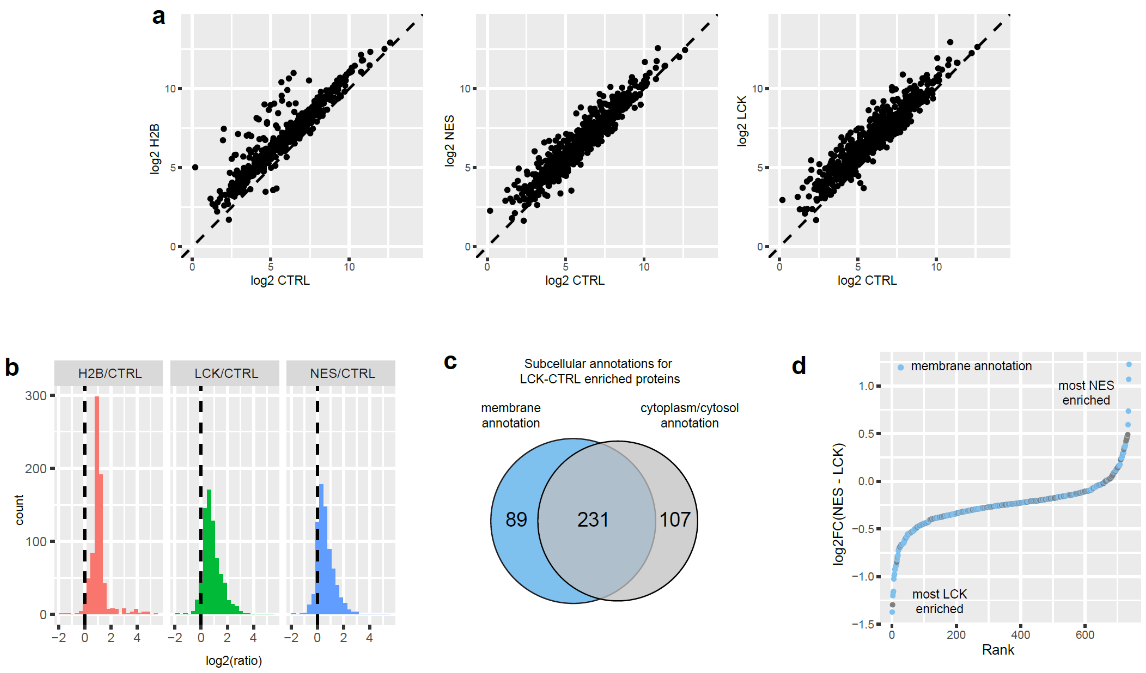


**Supplementary Figure 4. Extent of biotinylation measured by log2 ratio of protein intensities**

(a) Scatter plot between Cre-positive and Cre-negative samples. Mean of log2 protein intensities were used in the plot.

(b) Histogram of log2 ratio. Log2 ratio between Cre-positive and Cre-negative samples. Vertical dotted line, log2(ratio = 1) = 0, indicating no enrichment.

(c) Subcellular annotation for LCK-CTRL enriched proteins. Venn diagram showed that majority of LCK-CTRL enriched proteins have both membrane and cytoplasm annotations.

(d) Log2 ratio v.s. rank plot for NES – LCK. Lower log2 (NES – LCK) ratios indicated a greater enrichment by LCK construct.
